# Impaired Social Learning Predicts Reduced Real-life Motivation in Individuals with Depression: A Computational fMRI Study

**DOI:** 10.1101/652776

**Authors:** Anna-Lena Frey, Ciara McCabe

## Abstract

**Background:** Major depressive disorder is associated with altered social functioning and impaired learning, on both the behavioural and the neural level. These deficits are likely related, considering that successful social interactions require learning to predict other people’s emotional responses. Yet, there is little research examining this relation.

**Methods:** Forty-three individuals with high (HD; N=21) and low (LD; N=22) depression scores answered questions regarding their real-life social experiences and performed a social learning task during fMRI scanning. As part of the task, subjects learned associations between name cues and rewarding (happy faces) or aversive (fearful faces) social outcomes. Using computational modelling, behavioural and neural correlates of social learning were examined and related to real-life social experiences.

**Results:** HD participants reported reduced motivation to engage in real-life social activities and demonstrated elevated uncertainty about social outcomes in the task. Moreover, HD subjects displayed altered encoding of social reward predictions in the insula, temporal lobe and parietal lobe. Interestingly, across all subjects, higher task uncertainty and reduced parietal prediction encoding were associated with decreased motivation to engage in real-life social activities.

**Limitations:** The size of the included sample was relatively small. The results should thus be regarded as preliminary and replications in larger samples are called for.

**Conclusion:** Taken together, our findings suggest that reduced learning from social outcomes may impair depressed individuals’ ability to predict other people’s responses in real life, which renders social situations uncertain. This uncertainty, in turn, may contribute to reduced social engagement (motivation) in depression.

## 1 Introduction

Deficits in social functioning are commonly observed in major depressive disorder (MDD; Katz, Conway, Hammen, Brennan, & Najman, 2011; Rhebergen et al., 2010; Rottenberg & Gotlib, 2008). Compared to controls, depressed individuals have fewer friends (Brim et al., 1982; Frey et al., 2019; Youngren and Lewinsohn, 1980), fewer intimate relationships (Gotlib and Lee, 1989), and spend less time with people in their social circle (Youngren and Lewinsohn, 1980). Additionally, depressed subjects show inappropriate behaviour during social interactions (reviewed in Rottenberg & Gotlib, 2008; Segrin, 2000), which can result in the receipt of negative feedback from other people (Segrin and Abramson, 1994).

Successful interpersonal interactions require learning to predict other people’s responses and adjusting one’s own behaviour accordingly. Therefore, social functioning abnormalities in MDD may partly be linked to impaired learning from interpersonal outcomes. In line with this suggestion, we previously found that subjects with depression symptoms show deficits in learning from social feedback and demonstrate heightened negative feedback expectancy biases during a social decision-making task. Interestingly, impaired learning predicted the experience of more negatively perceived social encounters in real life, while negative biases, as well as social anhedonia, were associated with decreased amounts of time spent with friends (Frey et al., 2019). Moreover, using a social conditioning paradigm it has previously been observed that elevated depression scores are correlated with heightened arousal ratings in response to faces that had been paired with negative statements about the participant. This effect was still seen three months after the conditioning phase, indicating that the learning of negative social associations may be enhanced in individuals with higher levels of depressive symptomatology (Wiggert et al., 2017).

The above research provides limited evidence for changes in social learning in depressed individuals. Additionally, a range of studies have reported alterations in *non-social* learning in MDD. For instance, using decision-making tasks, it has been observed that depressed subjects display impaired reward learning (Blanco et al., 2013; Cooper et al., 2014; Herzallah et al., 2013; Kumar et al., 2018; Kunisato et al., 2012; Maddox et al., 2012; Pechtel et al., 2013; Robinson et al., 2012), while their punishment learning is either enhanced (Beevers et al., 2013; Maddox et al., 2012) or unchanged (Herzallah et al., 2013; Kumar et al., 2018; Kunisato et al., 2012; Robinson et al., 2012), when compared to controls. Moreover, in Pavlovian conditioning paradigms, depressed participants tend to demonstrate less accurate reward contingency predictions during or after the conditioning phase (Kumar et al., 2008; Robinson et al., 2012, although see Lawson et al., 2017 and Rupprechter, Stankevicius, Huys, Steele, & Seriès, 2018 for no group differences). By contrast, behavioural punishment conditioning does not seem to differ between depressed and control subjects when assessed with explicit measures (although neural group effects have been observed, see below; Lawson et al., 2017; Robinson et al., 2012).

The above behavioural research has been extended by neuroimaging studies which have examined neural learning signals with the use of computational models. In these models, the predictive value of a given cue is iteratively updated based on the difference between current outcomes and previous predictions. The latter difference, referred to as a prediction error (PE), as well as model-derived prediction values, have been used as parametric modulators in fMRI analyses.

Using this approach, it has been found that depressed individuals display reduced reward PE encoding in the midbrain, striatum, medial orbitofrontal cortex, dorsal anterior cingulate cortex, and hippocampus, compared to controls (Gradin et al., 2011; Kumar et al., 2018, 2008; Rothkirch et al., 2017). Interestingly, the magnitude of the striatal reward PE signal has been shown to moderate the relationship between real-life anticipatory and consummatory pleasure in depressed subjects (Bakker et al., 2018). Moreover, while some studies have observed attenuated habenula punishment PE representations in depression (Liu et al., 2017), others have found these representations to be unchanged in MDD (Rothkirch et al., 2017).

In addition, examinations of neural prediction encoding have found that depressed subjects display reduced reward prediction-related responses in the hippocampus and parahippocampus (Gradin et al., 2011), as well as decreased inverse correlations between reward prediction and PE signals in the ventral striatum (Greenberg et al., 2015), compared to controls. Additionally, depressed patients demonstrate reduced punishment prediction encoding in the habenula (when shocks are used as outcomes; Lawson et al., 2017).

The above findings suggest that depression is associated with learning deficits, both on the behavioural and the neural level, partly due to impaired generation and updating of outcome predictions. However, it should be noted that most previous studies assessing learning in MDD utilised non-social outcomes. Given the ubiquity of social stimuli in everyday life, it is important to further examine how far depressed subjects’ learning impairments extend to the social domain, and whether these impairments are related to the abovementioned social functioning deficits in MDD. The current study aimed to address this question. For this purpose, a social learning task was developed in which name cues were presented followed by faces that probabilistically displayed happy, neutral, or fearful expressions. Participants with high and low depression scores completed the task during fMRI scanning and were asked to learn the average likelihood of seeing a particular emotional expression after a given name cue. Additionally, subjects answered a number of questions about their real-life social experiences. A computational model was applied to the learning task data and model-derived prediction and PE values were used as parametric modulators in the fMRI analysis to assess the neural correlates of social learning. It was hypothesised that individuals with high depression scores would show impairments in the behavioural and neural prediction of social outcomes and that these deficits would be related to deficits in real-life social experiences.

## 2 Methods

### 2.1 Participants

The current study included 43 right-handed volunteers between the age of 18 and 45 years who scored below 8 (LD; N = 21) or above 16 (HD; N = 22) on the Beck Depression Inventory (BDI, Beck, Steer, & Brown, 1996). Subjects were screened using the structured clinical interview for DSM-IV (SCID; First, Spitzer, Gibbon, & Williams, 1996). LD volunteers were excluded if they had a history of any Axis I disorder or had ever taken any psychiatric medication. HD subjects were ineligible if they had ever experienced any Axis I disorder, apart from depression and moderate levels of secondary anxiety symptoms, or if they had taken any psychiatric medication in the past year. Additional exclusion criteria for volunteers in either group were the current use of any medications besides contraceptives, the use of recreational drugs in the past three months, smoking more than five cigarettes per week, or demonstrating contraindications to MRI scanning.

The study received ethical approval from the University of Reading Ethics Committee (UREC-16/08) and was carried out in accordance with the Declaration of Helsinki. All subjects provided informed consent.

### 2.2 Procedure

Before the testing session, potential participants attended a screening visit during which the SCID, as well as an interview about past and current medical conditions, were conducted to ascertain that none of the exclusion criteria were met. Subsequently, eligible subjects completed the following online questionnaires at home: trait subscale of the State and Trait Anxiety Inventory (STAI; Spielberger, Gorsuch, Lushene, Vagg, & Jacobs, 1983), Revised Social Anhedonia Scale (RSAS, Eckblad, Chapman, Chapman, & Mishlove, 1982), Uncertainty Intolerance Scale (UIS, Buhr & Dugas, 2002), and a demographics form.

In addition, subjects answered several questions about their everyday social interactions, indicating how many friends they have, how close they feel to these friends, and how difficult they find it to make new friends. Participants also rated their anticipatory, motivational and consummatory responses to pleasant social and non-social activities.

After the above questionnaires had been completed, a testing session was arranged. At the beginning of the session, participants filled in the Positive and Negative Affect Scale (PANAS; Watson, Clark, & Tellegen, 1988). Subsequently, they performed a name learning test (see supplement) and some practice trials of the social learning task outside the MRI scanner. Following the practice, subjects completed the social learning task in the MRI scanner, and, after the scan, filled in a task feedback questionnaire.

### 2.3 Social Learning Task

During the social learning task, participants’ aim was to learn how likely it is that a given name cue is followed by a happy, neutral or fearful facial expression. At the beginning of each trial, subjects saw one of the six names (1000ms), followed by a visual analogue rating scale (5000ms; see below). Subsequently, the face associated with the name was displayed (1000ms), showing either a neutral or an emotional expression, as determined by the probabilistic contingencies described below. The stimulus presentation was separated by a 2000ms inter-stimulus interval, and the inter-trial interval was jittered by drawing from an exponential distribution with a minimum of 2000ms and a mean of 2500ms (see Figure 1).

**Figure 1:**
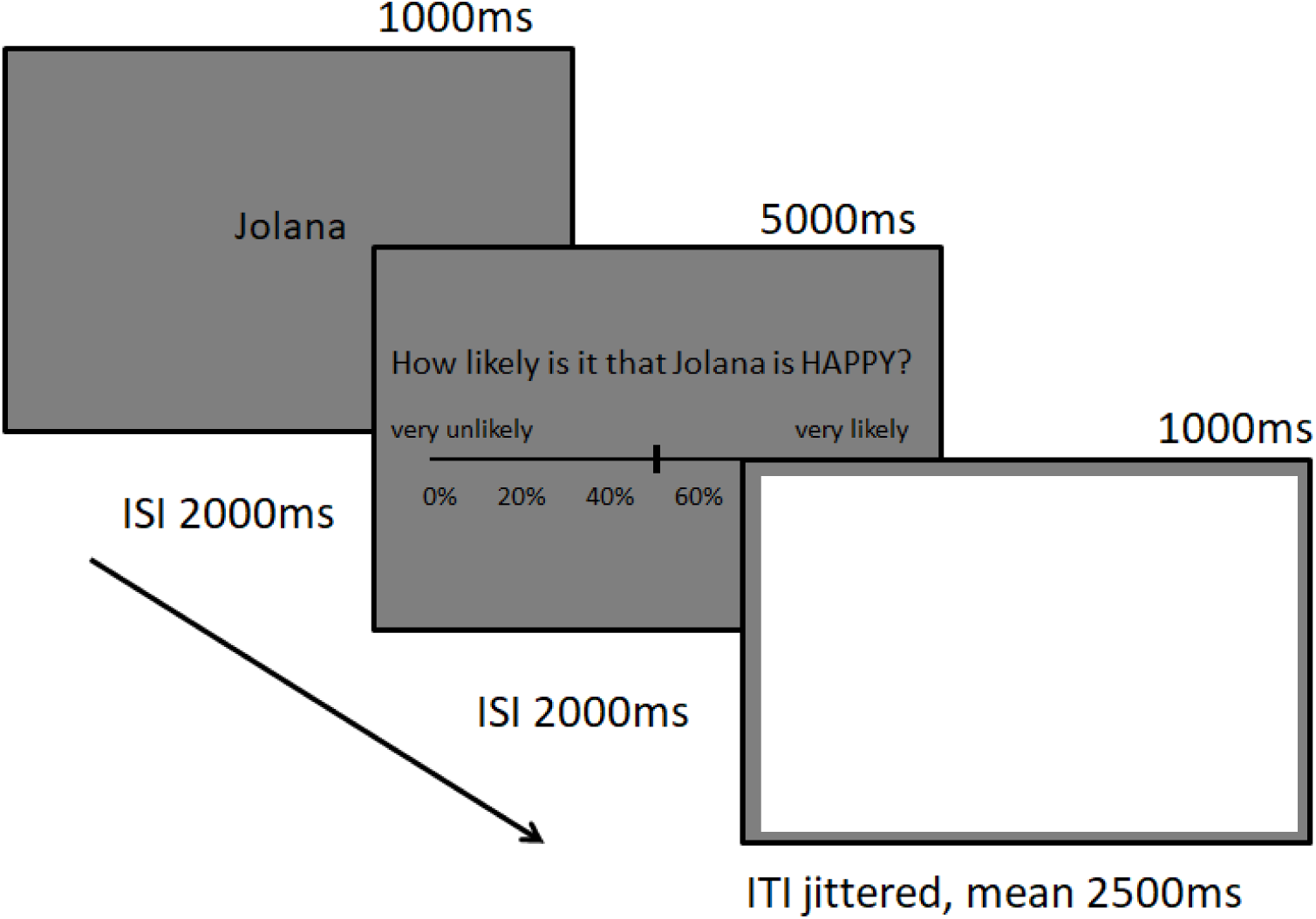
Example of a social learning task trial.

The task was divided into social reward and social aversion blocks which were performed in counterbalanced order. In the social reward block, three of the six faces were displayed, each of which had a different likelihood (25%, 50% or 75%) of showing a happy rather than a neutral expression. In the social aversion block, the other three faces were presented, each of which had a different likelihood (25%, 50% or 75%) of displaying a fearful rather than a neutral expression. The six faces were randomly assigned to the blocks and likelihoods for each participant and were presented in a pseudo-random order.

Subjects were asked to learn how likely it was, on average, that a given face displayed an emotional expression. They indicated this likelihood on a visual analogue scale, ranging from 0% to 100%, in response to the question ‘How likely is it that [name] is [HAPPY / AFRAID]?’. Participants were instructed to start with a guess, and to subsequently base their ratings on the intuition or ‘gut feeling’ they derived from all the times they had seen the name-face pairing before.

The task practice consisted of 8 repetitions of each name-face pairing, resulting in 24 trials per block and 48 practice trials in total (which were performed outside the MRI scanner). The experimental phase (which was completed inside the MRI scanner) included 12 presentations of each name-face pairing, resulting in 36 trials per block and 72 experimental trials in total.

### 2.4 Analysis

#### 2.4.1 Behavioural Analysis

Normality assumptions were not met for the questionnaire or name learning data. Group differences in these measures were therefore assessed using Mann-Whitney U tests.

Social learning task performance was examined by performing a mixed-measure (group x valence x probability) ANOVA on the likelihood ratings which were averaged across practice and experimental trials.

Moreover, to examine subjects’ uncertainty regarding the task outcomes, likelihood ratings were converted into uncertainty scores. For this purpose, 50 (i.e. the value indicating maximal uncertainty) was subtracted from each likelihood rating of a given participant, separately for social reward and aversion blocks. The resulting values were transformed into absolutes and then averaged across probabilities (separately for the two blocks). This yielded two scores for each subject, with lower scores indicating higher uncertainty about what outcomes to expect. To make the result interpretation more intuitive, scores were reversed by subtracting each score from the maximum value across all participants. Thus, in the below analysis high levels of uncertainty are indicated by high uncertainty scores. A mixed-measure (group x valence) ANOVA was performed on these scores.

Additionally, to relate the learning task performance to real-life measures, uncertainty scores were entered into a regression analysis. Given that the scores for social reward and aversion blocks were highly correlated (*r* = 0.57; *p* < 0.001), scores were averaged across the two blocks. The averaged uncertainty score was then mean-centred and used to predict participants’ motivation to engage in real-life social activities, together with BDI, RSAS, and mean-centred UIS negativity scores (calculated based on Sexton & Douglas 2009). An uncertainty score*UIS negativity interaction term was also included in the analysis, as it is likely that uncertainty about social outcomes primarily affects social engagement motivation when uncertainty is perceived as negative. STAI scores were not entered into the analysis, because this would have resulted in a violation of the multicollinearity assumption (Variance Inflation Factor > 10) due to a high correlation between STAI and BDI scores. This high correlation is in line with previous findings demonstrating that the STAI contains many items that map onto depression rather than specifically onto anxiety (Bados et al., 2010). However, it should be noted that STAI scores did not significantly contribute to the prediction of motivation when they were included in the regression model and BDI scores were removed.

#### 2.4.2 Computational Modelling

A standard Rescorla-Wagner model (Rescorla and Wagner, 1972) with a free learning rate parameter (α) was applied to the data (see supplement for details). Parameters were estimated by minimising the sum of squared errors between the model prediction value (multiplied by 100) and the participants’ likelihood ratings (similar to Hindi Attar, Finckh, & Büchel, 2012). Model parameter values and fits were compared between groups using Mann-Whitney U tests.

#### 2.4.3 fMRI Analysis

Functional MRI images were acquired using a three-Tesla Siemens scanner (Siemens AG, Erlangen, Germany) and the preprocessing and analysis of the data were performed using the Statistical Parametric Mapping software (SPM12; http://www.fil.ion.ucl.ac.uk/spm; see supplement for details). A first-level GLM analysis was conducted to examine the neural encoding of social outcome predictions. For this purpose, computational model-derived prediction values were entered as parametric modulators at the time of the cue, using separate regressors for the social reward and aversion blocks. On the second level, whole-brain one-way ANOVAs were conducted for group comparisons. Moreover, to relate the fMRI results to real-life measures, parameter estimates were extracted from the peak voxels of the prediction-related group contrast and were correlated with participants’ reported motivation to engage in positive social activities (similar to Gradin et al., 2011).

Additionally, neural prediction error (PE) encoding was examined. Brain responses encoding a canonical PE should, at the time of the outcome, covary positively with outcome values and negatively with prediction values. As in previous studies (e.g. Chowdhury et al., 2013; Rothkirch et al., 2017; Rutledge et al., 2017), these two PE components were thus entered into the first-level analysis as separate parametric modulators at the time of the outcome (for the social reward and social aversion block). Subsequently, MarsBar (Brett et al., 2002) was used to extract average parameter estimates for the two components from a 6mm sphere around striatal coordinates that have been found to encode PEs in a previous meta-analysis (left ROI: −10 8 −6; right ROI: 10 8 −10; Chase et al., 2015). The extracted values were then compared between groups using one-way ANOVAs.

## 3 Results

### 3.1 Behavioural Results

#### 3.1.1 Demographic and Questionnaire Measures

Mann-Whitney U tests revealed that there were no significant group differences in age (*U* = 219, *p* = 0.970). As expected, BDI (*U* = 0, *p* < 0.001), RSAS (*U* = 22, *p* < 0.001), STAI-T (*U* = 0, *p* < 0.001), UIS negativity (*U* = 17, *p* < 0.001), and PANAS Negative Affect Scale (*U* = 65, *p* < 0.001) scores were significantly higher in HD than in LD participants. Additionally, PANAS Positive Affect Scale scores were significantly lower in HD than in LD subjects (*U* = 349, *p* = 0.001; see Table 1).

**Table 1:** Demographic data and questionnaire scores for individuals with high (HD) and low (LD) depression scores.

|  | HD (N = 21) |  | LD (N = 22) |  |
| --- | --- | --- | --- | --- |
|  | Mean | SD | Mean | SD |
| Age (years) | 23.20 | 5.66 | 22.45 | 4.35 |
| N females/ males | 17/4 | - | 14/8 | - |
| BDI* | 26.05 | 9.63 | 1.36 | 1.84 |
| RSAS* | 18.57 | 6.43 | 5.77 | 4.31 |
| STAI-T* | 57.75 | 7.12 | 27.85 | 6.92 |
| UIS - neg* | 94.71 | 17.81 | 52.76 | 17.19 |
| PANAS - pos* | 24.38 | 5.71 | 31.52 | 6.57 |
| PANAS - neg* | 21.29 | 7.27 | 13.43 | 5.26 |

SD, standard deviation; BDI, Beck Depression Inventory; RSAS, Revised Social Anhedonia Scale; STAI-T, trait score of the State Trait Anxiety Inventory; UIS - neg, Uncertainty Intolerance Negativity Scale; PANAS-pos/neg, positive and negative mood scores of the Positive and Negative Affect Scale;
\* asterisks indicate significant group differences

#### 3.1.2 Real-Life Social Experiences

Compared to LD subjects, HD participants indicated having significantly fewer friends (*U* = 320, *p* = 0.001), feeling less close to their friends (*U* = 364, *p* < 0.001), and finding it more difficult to form new friendships (*U* = 47, *p* < 0.001).

Moreover, HD individuals demonstrated significantly reduced motivation to engage in pleasant social activities (*U* = 294, *p* = 0.003), as well as significantly decreased anticipation (*U* = 316, *p* < 0.001) and enjoyment (*U* = 323, *p* < 0.001) of pleasant social activities, compared to LD controls. By contrast, no group differences were observed for anticipatory (*U* = 223, *p* = 0.365), motivational (*U* = 227, *p* = 0.309), or consummatory (*U* = 226, *p* = 0.322) responses to pleasant *non-social* activities.

#### 3.1.3 Social Learning Task Performance

A mixed measure ANOVA (group x valence x probability) performed on participants’ likelihood ratings revealed the expected main effect of probability (*F*(2, 82) = 94.95, *p* < 0.001), with participants rating the likelihood of seeing an emotional expression higher after cues that were more likely to be followed by an emotional face. Moreover, a main effect of valence was observed (*F*(1,41) = 8.30, *p* = 0.006) which indicated that participants rated the overall likelihood of seeing happy faces as higher than the likelihood of seeing fearful faces. Additionally, a group by probability interaction was found (*F*(2,82) = 11.77, *p* < 0.001) which was followed up as described below. All other main effects and interactions were not significant (all F < 2.3).

Follow-up one-way ANOVAs revealed that, compared to LD controls, HD participants’ likelihood ratings were significantly *lower* on trials with a 75% chance of showing a happy (*F*(1,41) = 9.12, *p* = 0.004) or fearful (*F*(1,41) = 3.98, *p* = 0.053) expression. By contrast, HD subjects’ ratings were significantly *higher* than those of controls on trials with a 25% chance of showing a happy (*F*(1,41) = 9.82, *p* = 0.003) or fearful (*F*(1,41) = 10.18, *p* = 0.003) face (see Figure 2). No group differences were found on trials with a 50% chance of displaying a happy (*F*(1,41) = 0.15, *p* = 0.698) or fearful (*F*(1,41) = 0.07, *p* = 0.796) expression.

**Figure 2:**
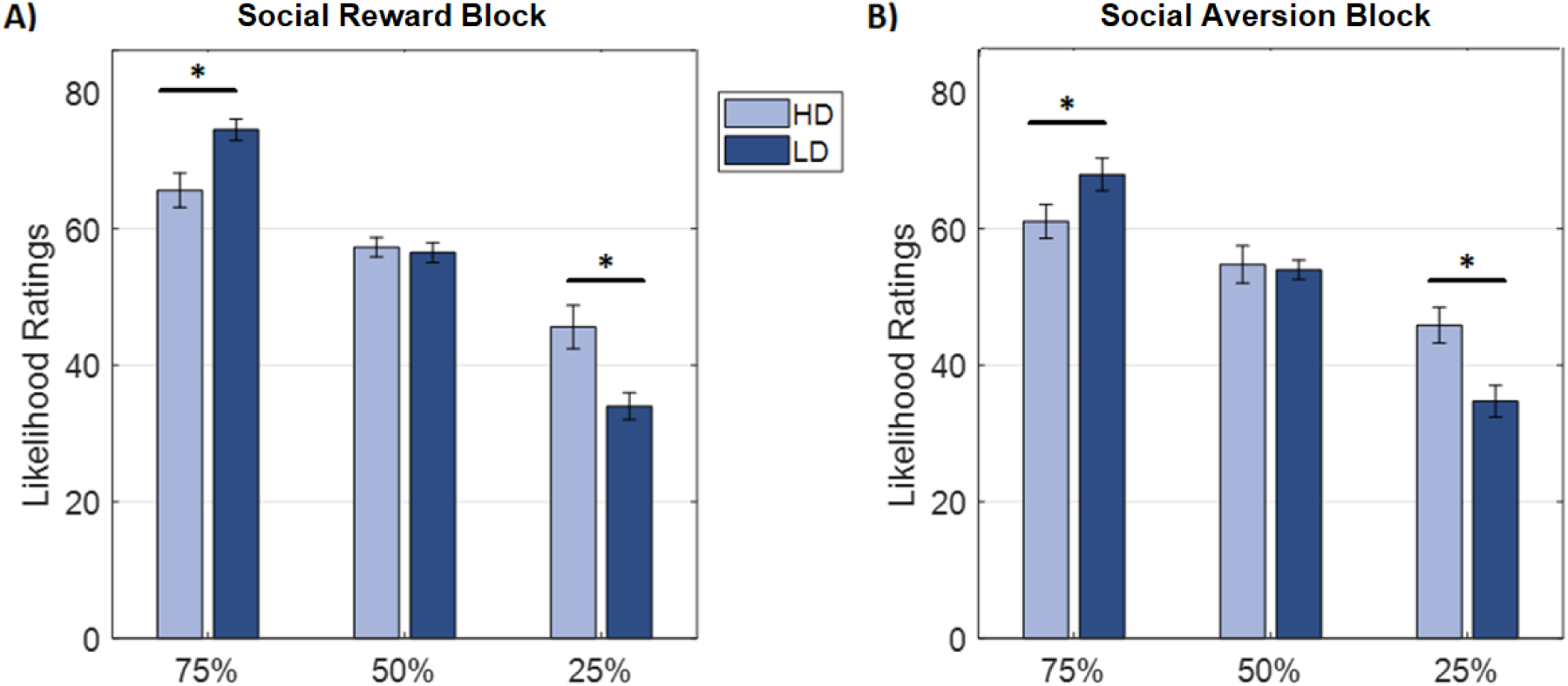
Likelihood ratings by chance of seeing an emotional face for A) the social reward and B) the social aversion block in individuals with high (HD) and low (LD) depression scores.

Moreover, a mixed-measure (group x valence) ANOVA conducted on participants’ uncertainty scores (which indicate the average difference between subjects’ ratings and 50%; see section 2.3.1) revealed a significant main effect of group, as HD subjects tended to be more uncertain about the social task outcomes than LD controls (*F*(1,41) = 3.67, *p* = 0.062). Additionally, a significant main effect of valence was found, showing that subjects were more uncertain about aversive than about rewarding outcomes (*F*(1,41) = 6.62, *p* = 0.014). No significant interaction effect was observed (*F*(1,41) = 0.160, *p* = 0.692).

Additionally, a multiple regression analysis revealed that task uncertainty scores (averaged across blocks), together with questionnaire measures, predicted participants’ motivation to engage in pleasant social activities (*F*(5, 32) = 8.57, *p* < 0.001, R^2^ = 0.51). Predictors significantly contributing to this relation were the main effect of UIS negativity (β = −0.55, *p* = 0.008), the UIS negativity * task uncertainty interaction term (β = −0.32, *p* = 0.015; see Figure 3), and, marginally, RSAS social anhedonia scores (β = −0.37, *p* = 0.061). By contrast, the main effect of task uncertainty (β = −0.21, *p* = 0.096) and BDI scores (β = 0.32, *p* = 0.149) had no significant effect. Thus, the motivation to engage in pleasant social activities was particularly reduced in individuals who were uncertain about what social outcomes to expect and who experienced uncertainty as negative.

**Figure 3:**
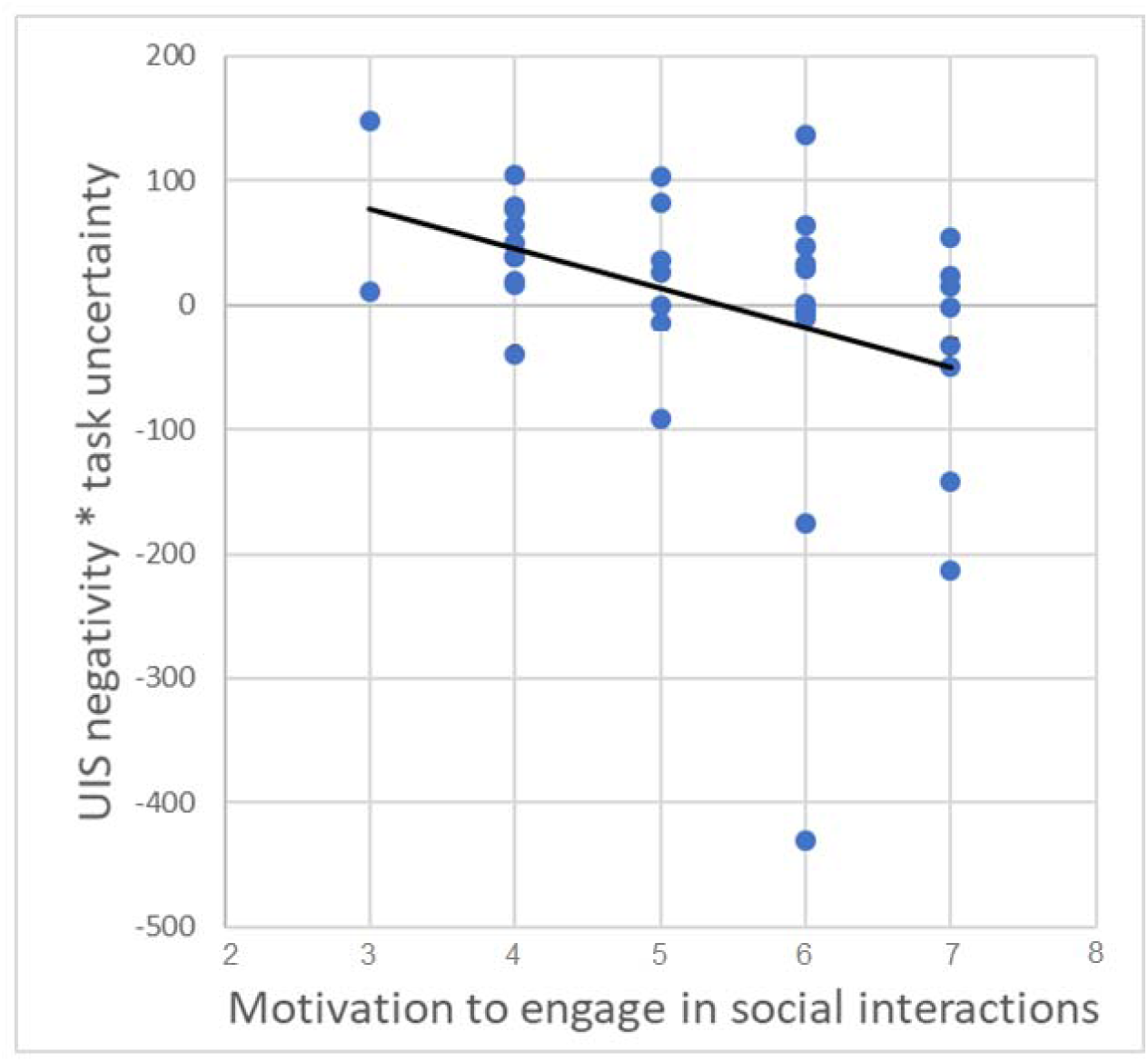
Scatter plot showing the association between motivation to engage in pleasant social activities (higher scores indicate higher motivation) and uncertainty intolerance (UIS) * task uncertainty interaction values.

#### 3.1.4 Computational Modelling

Mann-Whitney U tests on the model parameters revealed that learning rates were significantly lower in HD than in LD participants, both in the social reward (*U* = 351, *p* = 0.004) and in the social aversion (*U* = 355, *p* = 0.003) block. The model fit, as indicated by the sum of squared errors, did not differ significantly between groups in either the social reward (*U* = 171, *p* = 0.145; *U* = 169, *p* = 0.132) or aversion (*U* = 189, *p* = 0.308; *U* = 182, *p* = 0.234) block when using individual or averaged parameters (respectively).

### 3.2 fMRI Results

#### 3.2.1 Neural Prediction Value Encoding

Social reward (i.e. happy expression) prediction encoding was reduced in HD, compared to LD, subjects in the superior parietal lobe/ precuneus, as well as in a cluster including the right insula, supramarginal gyrus and superior temporal lobe (see Table 2 and Figure 4). No group differences were found for social aversion (i.e. fearful expression) prediction encoding.

**Figure 4:**
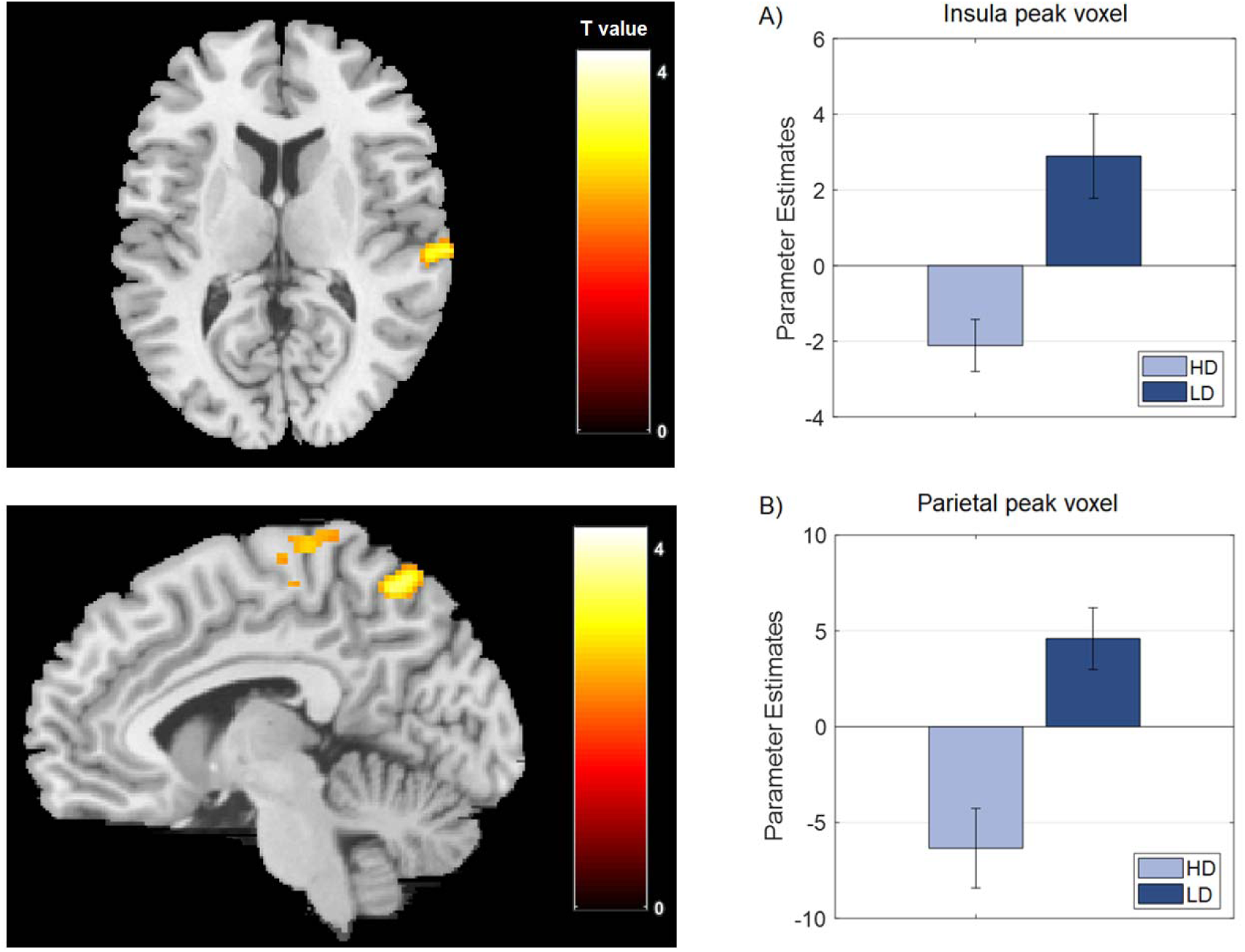
Clusters showing lower social reward prediction encoding in individuals with high (HD) than with low (LD) depression scores, as well as parameter estimates extracted from A) the insula peak voxel and B) the parietal peak voxel.

**Table 2.** Parametric modulation results for social reward prediction encoding in individuals with low (LD) vs. high (HD) depression scores.

|  | MNI coordinates |  |  |  |  |
| --- | --- | --- | --- | --- | --- |
| Brain Region | X | Y | Z | Z score | *p* value |
| LD > HD |  |  |  |  |  |
| Superior Parietal Lobe/ Precuneus | -18 | -58 | 68 | 3.80 | 0.001 |
| Right Insula | 48 | -20 | 18 | 3.47 | 0.045 |
| Right Supramarginal Gyrus | 58 | -32 | 24 | 3.28 |  |
| Right Superior Temporal Lobe | 68 | -22 | 12 | 3.17 |  |

Whole-brain cluster p values are family-wise error corrected at $p < .05$

Across all subjects, correlation analyses revealed a significant positive correlation between participants’ motivation to engage in pleasant social activities and parameter estimates extracted from the peak prediction-related group comparison voxels in the parietal lobe (*r* = 0.49, *p* = 0.002) and insula (*r* = 0.36, *p* = 0.023). This relationship remained significant for the parietal lobe (*r* = 0.36, *p* = 0.027), but not the insula (*r* = 0.25, *p* = 0.137), when BDI and task uncertainty scores were controlled for.

#### 3.2.2 Neural Prediction Error Encoding

One-way ANOVAs were conducted on the average parameter estimates extracted from a left and a right striatal ROI for the encoding of outcome and inverse prediction values (i.e. the two PE components). No significant group differences were found for either the social reward or the social aversion block (all F < 2.9).

## 4 Discussion

### 4.1 Uncertainty about social outcomes predicts reduced social engagement motivation

The current study examined learning from social outcomes in individuals with high (HD) and low (LD) depression symptoms, linking task performance to measures of real-life social experiences.

It was found that, in both the social reward and the social aversion block of our learning task, HD individuals *under*estimated the likelihood of being presented with emotional faces on *high* probability trials, while they *over*estimated this likelihood on *low* probability trials. In other words, HD subjects provided ratings close to 50% across all trial types, indicating general uncertainty about what outcomes to expect. These findings are partly consistent with previous reports of impaired reward conditioning in depression (Kumar et al., 2008; Robinson et al., 2012; see also Chen et al., 2015). Yet, it may seem somewhat surprising that HD subjects demonstrated higher uncertainty (and thus decreased learning) in the social aversion block, considering that past studies have observed *enhanced* punishment learning in depression (Beevers et al., 2013; Maddox et al., 2012). A possible explanation of this finding is that the social stimuli used in the current study may have been particularly likely to induce rumination in HD individuals, which may have interfered with the aversion learning process (Whitmer et al., 2012). Moreover, it is worth noting that, unlike previous tasks, the current paradigm required the *continuous* formation, updating and working memory maintenance of *explicit* outcome contingencies. This may have been particularly difficult for HD individuals (independent of the stimulus valence), which would explain the general learning deficit and increase in uncertainty observed in this group.

Notably, in everyday social cognition both implicit and explicit processes play a role (Frith and Frith, 2008). Thus, HD individuals’ impaired ability to explicitly predict other people’s responses is likely to have an effect on real-life social functioning. In line with this suggestion, the current study found that task-based uncertainty, in interaction with the perceived negativity of uncertainty, significantly predicted participants’ motivation to engage in positive social activities (even when depression scores were controlled for). That is to say, subjects who demonstrated more uncertainty about (and thus worse learning from) social outcomes in the task, and who were more averse to uncertainty in general, were less motivated to engage in pleasant social activities in real life. It is noteworthy that HD subjects demonstrated high levels of task uncertainty, regarded uncertainty as negative, and displayed reduced social engagement motivation. Taken together, these findings suggest that deficits in learning from social outcomes may contribute to social disengagement in depressed individuals. Social withdrawal, in turn, may further increase depressed subjects’ uncertainty regarding social encounters by reducing their exposure to situations in which social outcome contingencies can be learned.

The current findings are consistent with previous observations of increased intolerance of uncertainty in depression (Carleton et al., 2012). Moreover, past studies have reported a link between uncertainty intolerance and depressive rumination (Yook et al., 2010), and it has been argued that uncertainty leads to behavioural inhibition when it is regarded as negative (Carleton, 2016). It may thus be the case that, in response to higher social outcome uncertainty, depressed individuals are prone to ruminate about possible negative outcomes, which reduces (/inhibits) their motivation to engage in social activities. This idea is supported by the supplementary analysis of the present study which shows that the interaction between enhanced task uncertainty and *inhibitory* uncertainty intolerance predicts reduced social engagement motivation. In addition, the above suggestion is in line with our previous findings showing that increased negative social feedback expectancies are associated with social disengagement in individuals with high depressive symptomatology (Frey et al., 2019). It would be of interest for future studies to examine whether the relation between uncertainty and social disengagement is indeed mediated by rumination-induced negative expectancies.

### 4.2 Neural predication of social rewards is impaired in HD subjects

Consistent with the behavioural findings, the current study found that HD individuals displayed impaired learning signals on the neural level. Specifically, compared to controls, HD participants displayed lower covariation between social reward prediction values and BOLD responses in the superior parietal lobe, as well as in a cluster extending from the insula to the supramarginal gyrus and superior temporal lobe.

Given the superior parietal lobe’s involvement in attentional processing (Behrmann et al., 2004), this region may have been recruited because the repeated pairing of cues with happy expressions made the cues more salient targets for active attentional processing. Moreover, the insula, supramarginal gyrus and temporal lobe have previously been implicated in the processing (Fusar-Poli et al., 2009) and working memory maintenance (Nichols, Kao, Verfaellie, & Gabrieli, 2006) of faces. Hence, the increased engagement of these regions by cues that were more frequently paired with task-relevant happy expressions may reflect a working memory mechanism that aids the learning processes.

Based on the above, the findings of reduced social reward prediction encoding in HD individuals in the above regions could be taken to indicate a deficit in neural attention and working memory processing during learning. However, it should be noted that BOLD responses were not simply reduced in HD subjects, but were instead reversed. That is to say, rather than being close to zero, parameter estimates extracted from the peak voxels of the group contrast were significantly below zero in the HD group (and significantly above zero in the LD group; see supplement). This indicates that, in HD individuals, BOLD responses were higher the more frequently cues were associated with *neutral* faces. A possible explanation for this finding is that, due to negative processing biases, HD individuals perceived the ambiguous neutral faces as negative, especially when they were displayed amongst happy expressions. Such a negative perception may have made the neutral faces particularly salient, and may thus have led to the recruitment of attentional and working memory resources to represent and predict neutral rather than happy faces.

The above suggestion is consistent with previous behavioural observations showing that depressed individuals tend to perceive neutral expressions as negative (Bouhuys et al., 1999; Hale et al., 1998; Leppannen et al., 2004). Moreover, the increased salience of neutral faces may also have contributed to the behavioural findings of the current study. Specifically, the mismatch between task demands (of happy expression prediction) and neural processes (of neutral expressions prediction) may have given rise to the uncertainty reflected in HD participants’ task ratings. Notably, a similar mechanism could play a role in real life, if automatic processing supports learning from negative social feedback and reflective processes are needed (but potentially unable) to accurately predict the positive value of engaging in social activities (along the lines of the dual process model of Beevers, 2005).

It thus seems plausible that the neural processes of HD subjects may have supported the prediction of negatively perceived neutral expressions rather than that of happy faces. Following on from this suggestion, it may have been expected that the neural response to happy vs. neutral faces would have differed between groups, due to increased (aversive) processing of neutral faces in HD participants. Yet, such a group effect was not observed. This may potentially be the case because the prediction of neutral expressions in HD subjects, after some learning had occurred, may have engaged preparatory downregulation processes resulting in similar neural responses to neutral faces in HD and LD individuals.

Interestingly, the current study further found that lower social reward prediction encoding in the parietal lobe was significantly correlated with reduced motivation to engage in positive social activities in real life, even when task uncertainty and depression scores were controlled for. Considering the abovementioned involvement of the parietal lobe in attentional processing (Behrmann et al., 2004), this may indicate that individuals who demonstrate diminished attentional processing of positive social feedback, or enhanced attentional processing of ambiguous feedback, may be less motivated to engage in social activities (although the direction of this relation cannot be determined based on the present data). This may especially be the case in HD subjects, who displayed decreased parietal prediction encoding, as well as reduced motivation to engage in pleasant social situations. In line with this notion, we recently found that adolescents with depression symptoms displayed blunted anticipatory responses to reward in the precuneus (and insula) and showed reduced motivation (/effort) to gain rewards (Rzepa and McCabe, 2019).

### 4.3 Limitations

It should be noted that the current study included a relatively small sample size. Therefore, the results should be regarded as preliminary and replications in larger samples are called for. Moreover, it would be advisable for future studies to assess how social learning in depression is affected when other social stimuli (besides happy, neutral and fearful faces) are used.

### 4.4 Conclusion

All in all, the results of the current study indicate that individuals with high depression scores demonstrate impaired learning from social outcomes, on both the neural and the behavioural level. Importantly, this deficit was associated with reduced motivation to engage in real-life social activities, possibly due to increased negatively-perceived uncertainty about what to expect from social encounters. These findings tentatively suggest that improving social learning may contribute to reducing social withdrawal in depression. Future studies are needed to examine this suggestion.

## Acknowledgements

We would like to thank Evelyn Toh and Canan Asli Can for their assistance with the data collection.

